# Upregulating ANKHD1 in PS19 mice reduces Tau phosphorylation and mitigates Tau-toxicity-induced cognitive deficits

**DOI:** 10.1101/2024.11.15.623890

**Authors:** Xiaolin Tian, Nathan Le, Yuhai Zhao, Dina Alawamleh, Andrew Schwartz, Lauren Meyer, Elizabeth Helm, Chunlai Wu

## Abstract

Abnormal accumulation of Tau protein in the brain disrupts normal cellular function and leads to neuronal death linked with many neurodegenerative disorders such as Alzheimer’s disease. An attractive approach to mitigate Tau-induced neurodegeneration is to enhance the clearance of toxic Tau aggregates. We previously showed that upregulation of the fly gene *mask* protects against FUS- and Tau-induced photoreceptor degeneration in fly disease models. Here we have generated a transgenic mouse line carrying Cre-inducible ANKHD1, the mouse homolog of *mask*, to determine whether the protective role of *mask* is conserved from flies to mammals. Utilizing the TauP301S-PS19 mouse model for Tau-related dementia, we observed that ANKHD1 significantly reduced hyperphosphorylated human Tau in 6-month-old mice. Additionally, there was a notable trend towards reduced gliosis levels in these mice, suggesting a protective role of ANKHD1 against TauP301S-linked degeneration. Further analysis of 9-month-old mice revealed a similar trend of effects. Moreover, we found that ANKHD1 also suppresses the cognitive defect of 9-month-old PS19 female mice in novel object recognition (NOR) behavioral assay. Unlike previous therapeutic strategies that primarily focus on inhibiting Tau phosphorylation or directly clearing aggregates, this study highlights the novel role of ANKHD1 in promoting autophagy as a means to mitigate Tau pathology. This novel mechanism not only underscores ANKHD1’s potential as a unique therapeutic target for tauopathies but also provides new insights into autophagy-based interventions for neurodegenerative diseases.

## Introduction

Many neurodegenerative diseases are characterized and differentially diagnosed by intracellular deposits of insoluble proteins. The microtubule associated protein Tau is one of these proteins that form fibrillary aggregates in neurons and glial cells in patients with a variety of disorders collectively called tauopathies ^1,2^. As a microtubule-binding protein, Tau’s cellular function is to regulate microtubule dynamics. Indeed, more than 80% Tau is bound to microtubule (MT) under physiological conditions^3^. By interacting with MT and motor proteins such as Kinesin and Dynein, neuronal Tau proteins play an important role in stabilizing microtubules and cargo delivery from soma to presynaptic terminal through MT-mediated axonal transport^4^. However, in certain neurodegenerative disorders, Tau undergoes abnormal post-translational modifications and forms fibrillary aggregates in neurons and glial cells, namely paired helical filaments (PHF) of Tau proteins, which are twisted, rope-like structures formed by Tau proteins^1,5^. The formation of PHF of Tau proteins (PHF-Tau) is a hallmark and considered to be one of the direct causal factors for diseases collectively known as tauopathies, such as Alzheimer’s disease, frontotemporal dementia, and corticobasal degeneration. These toxic PHF-Tau aggregates could spread like prions within a cell and across cell-cell boundary as shown in both cell cultures^6,7^ and in mice^8^.

Emerging evidence indicates that early Tau mislocalization, oligomerization, and solubility alterations are more closely linked to toxicity than the later stages of highly ordered tau filaments^1,9–11^. Therefore, targeting tau clearance at pre-symptom stage offers a promising therapeutic approach.

Autophagy is crucial for the removal of aggregated proteins and serves as a primary clearance route for tau in healthy neurons. Impairment of the autophagy-lysosomal pathway has been identified in the brains of tauopathy patients^12–14^, as well as in animal models^15^. In the mouse models, autophagy activators have been shown to reduce levels of misfolded and aggregated proteins, thereby mitigating tau spread and the related neuronal loss, highlighting the therapeutic potential of autophagy modulators^15–17^. In *Drosophila*, we recently showed that Mask is necessary and sufficient for promoting the autophagy-lysosomal pathway^18^. In the fly model of Tauopathy, upregulation of Mask in the fly photoreceptors suppresses, while downregulation of Mask enhances, the eye degeneration induced by expressing human mutant Tau in the photoreceptors^18^. These data are consistent with the notion that Mask-mediated increase of autophagy pathway plays a cytoprotective role against degeneration induced by toxic protein aggregates in flies.

Mammals express two *mask* homologues, *ANKRD17* and *ANKHD1*. Loss of function of *ANKRD17* has been linked to neural developmental disorders^19–21^, while the levels of ANKHD1 are elevated in numerous tumors^22–27^. Studies into the role of ANKHD1 in cancers demonstrate a crucial role in driving uncontrolled cellular proliferation, enhanced tumorigenicity, cell cycle progression, and increased epithelial-to-mesenchymal transition, resulting in greater tumor infiltration, increased metastasis, and larger tumours^23,27–31^. ANKHD1 comprises 25 ankyrin motifs organized into two domains, along with a single KH domain. When compared to Mask, the first set of ankyrin domains shows a 79% sequence similarity, the second set has a 63% sequence similarity, and the KH domain exhibits a 54% sequence similarity. Given ANKHD1’s pro-survival function cancer cells and its high sequence homology with Mask, We hypothesize that ANKHD1 exerts a conserved neuroprotective role similar to its Drosophila homolog Mask, by promoting autophagy, which in turn reduces pathological Tau accumulation and alleviates Tau-induced neurotoxicity. Our data show that expression of ANKHD1 in neurons in the mouse transgenic model that expresses exogenous human mutant P301S Tau ameliorated the accumulation of hyper-phosphorylated Tau, reduced gliosis in the same hippocampal region and partially restored the cognitive deficits caused by the mutant Tau protein.

## Results

### Generation and validation of a Cre-inducible ANKHD1 expression construct

We have generated a Cre-inducible ANKHD1 expression construct under the CMV promoter (Fig. 1A). The design of this construct allows the Cre proteins to remove the *loxp*-flanked GFP STOP cassette, and therefore, permits the expression of ANKHD1 as well as the mCherry protein whose coding sequence follows ANKHD1 and an IRES signal (Fig. 1). Before this Cre-inducible ANKHD1construct was used to generate transgenic mouse colonies, we transfected HEK293 cells with the construct along with a CMV-Cre plasmid. As anticipated, in the absence of CMV-Cre, we observed most of the cells strongly expressed GFP (Fig. 1B left). In the presence of CMV-Cre, GFP expression was largely diminished, and modest mCherry expression was detected (Fig. 1B right). We predicted that mCherry-expressing cells also express ANKHD1, which was confirmed by western blot using anti-ANKHD1 antibody (Fig. 1C).

**Figure 1.**
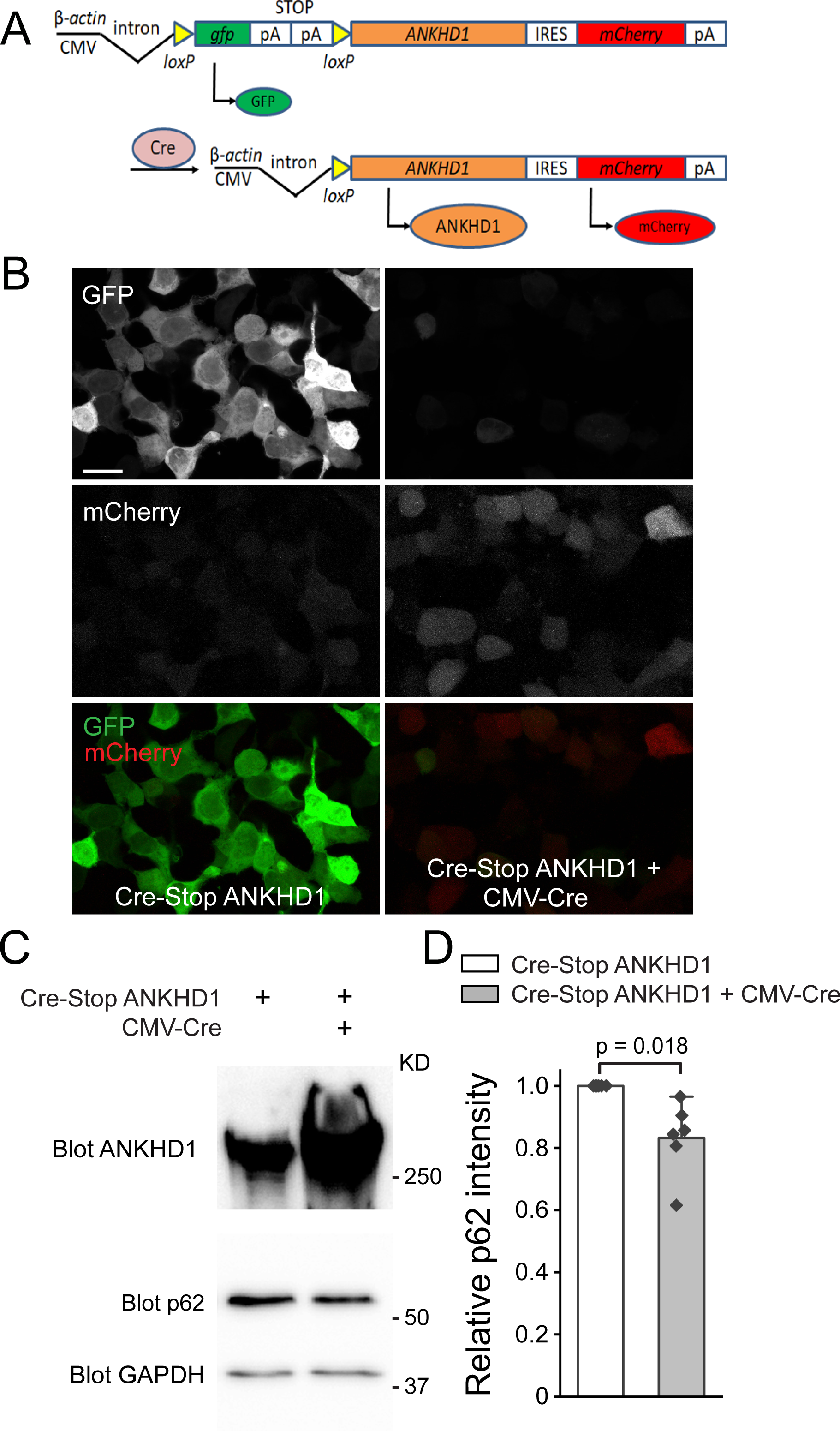
Generating Cre-inducible ANKHD1 transgenic mouse. (**A**) Schematics of the DNA construct for generating Cre-inducible ANKHD1 mouse. (**B**) Representative confocal images of GFP and mCherry auto-fluorescence from HEK293 cells transfect with Cre-Stop ANKHD1 with (right) or without (left) CMV-Cre plasmid. Scale bar, 20 μm (**C**) Western blots of cell lysates from the two groups of HEK293 cells showing in B with anti-ANKHD1, p62, and GAPDH antibodies (as loading control). (**D**) Quantification of p62 intensity normalized to GAPDH.

To assess the impact of ANKHD1 on autophagy flux, we measured the levels of p62, a well-established marker of autophagy activity. Under normal conditions, p62 facilitates the transport of damaged cellular components to autophagosomes and is subsequently degraded in the autolysosomes. We observed significant reduction of p62 protein levels in ANKHD1 expressing HEK293 cells (Fig. 1 CD), suggesting that ANKHD1 overexpression elevates autophagy flux.

This finding aligns with our hypothesis that ANKHD1 promotes autophagy and supports its potential role in mitigating Tau pathology.

### Generation and validation of Cre-inducible ANKHD1 expression mouse lines

The Cre-inducible ANKHD1 constructs were then used to generate transgenic mice. We obtained three founder lines, #38, #48, and #62. Founder line #38 was chosen for the rest of this study based on its consistent genotyping by PCR analysis (Supplemental Fig. 1).

ANKHD1 suppresses neuron-specific histopathological lesions induced by mutant human Tau proteins in the mouse brain.

To test whether ANKHD1 is capable of conferring a conserved protective effect in neurons, we crossed the Cre-inducible ANKHD1 transgene into the previously established TauPS19, or pnp- Tau-P301S mouse model^32^, and examined the neuropathological hallmarks in the mouse brains and mouse behaviors that reflect their cognitive functions. We generated a cohort of mice littermates that contains the control mice (+/+; CamK2a-Cre/+), the Tau^PS19^ mice (pnp-Tau- P301S/+; CamK2a-Cre/+), and Tau^PS19^-ANKHD1 mice (Cre-Stop-ANKHD1/+; pnp-Tau- P301S/+; CamK2a-Cre/+) through two sequential crosses: the first cross uses the TauPS19 and the CamK2a-Cre as the parental mice; and the second cross uses the resulting pnp-Tau-P301S/+; CamK2a-Cre/+ progenies from the first cross to mate with the Cre-inducible ANKHD1 transgenic mice.

Previous characterization of the TauPS19 mice has already shown that Tau-linked aggregates and related cellular deficits manifested as early as 3-month-old age and became prominent at 6- month-old age, although neuronal loss in hippocampus and cognitive decline were not evident at these ages^32^. We then examined the levels of Tau hyperphosphorylation and gliosis in the hippocampal region in the mice brains at the 6-month age. As expected, we observed prominent levels of PHF-Tau in the hippocampal CA3 region of the brain slices from Tau^PS19^ mice (pnp-

Tau-P301S/+; CamK2a-Cre/+) marked by the reactivity to the AT8 antibody that specifically recognizes the hyperphosphorylated Tau protein. Interestingly, in the brain slices of the Tau^PS19^- ANKHD1 mice, we detected significant decrease of the phosphorylated Tau levels in the same brain region (36% reduction in the male and a 42% reduction in the female), suggesting that ANKHD1 is able to mitigate the accumulation of phosphorylated Tau aggregates in the mouse brain (Fig. 2).

**Figure 2.**
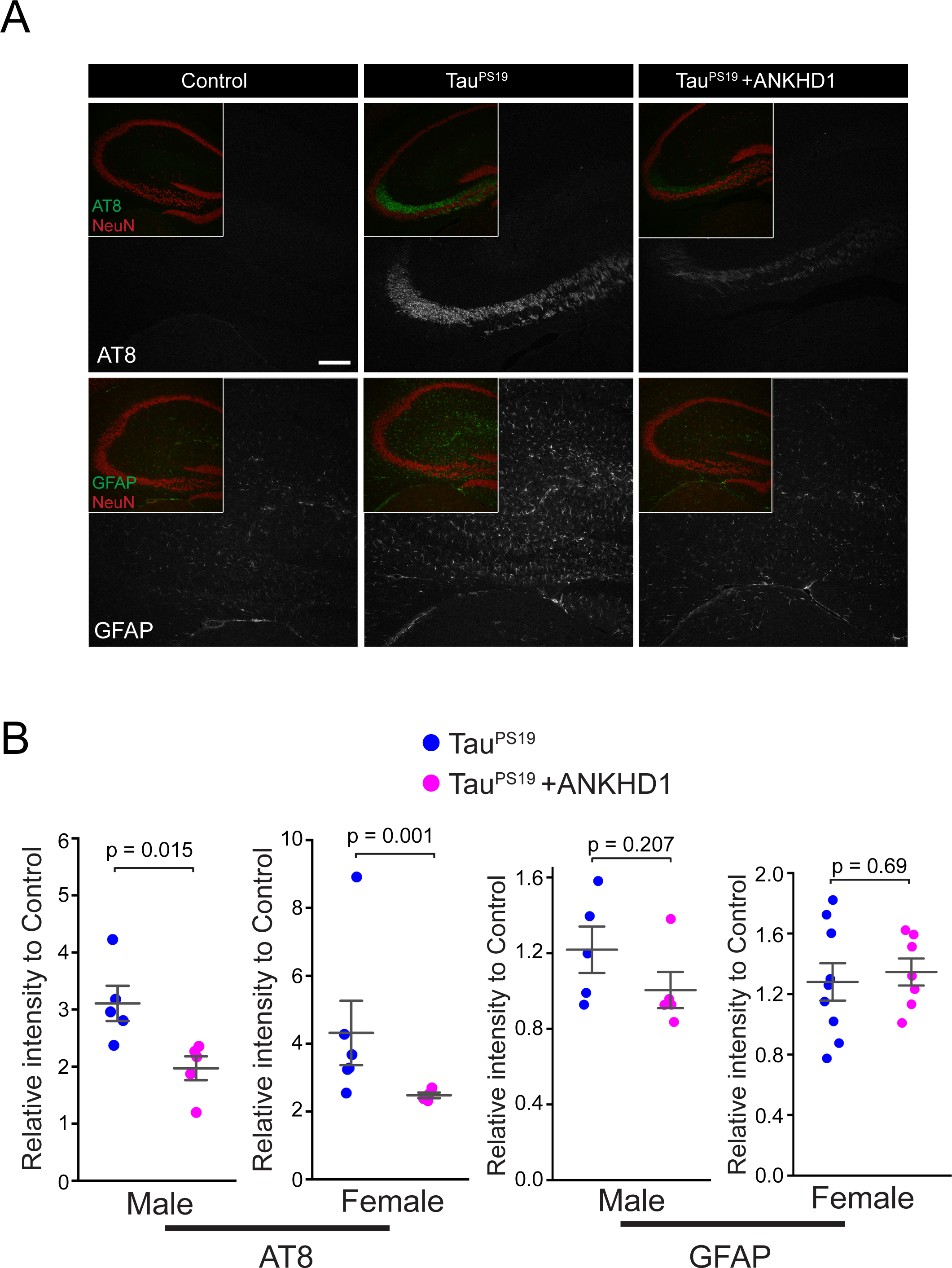
ANKHD1 co-expression reduces phosphorylated Tau in the brain of TauP301S transgenic mice at 6-month of age. (**A**) Representative confocal images of mouse brain hippocampus of Control, Tau^P301S^, and Tau^P301S^ + ANKHD1 mice. Mouse brain sections were stained with AT8 (anti-Tau-phospho-Ser202 and phosho-Thr205), anti-GFAP, and anti-NeuN antibodies, respectively, to detect hyperphosphorylated Tau, microglial cells, and hippocampal areas. (**B**) AT8 and GFAP intensities in the brain sections from both male and female mice were quantified. Average fluorescence intensity of AT8 or anti-GFAP was normalized to their control littermates. The data were presented as the relative intensity ratio.

Gliosis was a concomitant histopathological sign to neuronal loss in tauopathies^33^. In the PS19 transgenic mouse, weak increase of glial fibrillary acidic protein (GFAP), a marker for astroglia cells, could be detected throughout brain at 3-month-old age, while marked increase of GFAP was observed in 6 month old PS19 mice^32^. We then examined the ability of ANKHD1 to suppress glial activation caused by the mutant Tau protein in hippocampus. By comparing the TauPS19-ANKHD1 mice to the TauPS19 mice, we observed a trend of reduced gliosis levels in the male (17% reduction), but not in female, littermates that carry the ANKHD1 transgene (Fig. 2).

We next continued the histological analysis with the 9-month-old mice using same assays. We observed similar trends of reduced hyperphosphorylated Tau (17% reduction) and GFAP (9% reduction) intensities in the TauPS19-ANKHD1 male mice. (Fig. 3). The TauPS19 mice exhibit neuronal loss in hippocampus at 9-month-old age^32^, and in the same mice cohort, we also observed a trend of preserved NeuN staining (23% increase) in hippocampus in the brain slices of the TauPS19-ANKHD1 mice compared to the TauPS19 mice (Fig. 3). Together, ANKHD1 shows beneficial effects to reduce the accumulation of phosphorylated Tau, and related neuronal loss and gliosis in hippocampus.

**Figure 3.**
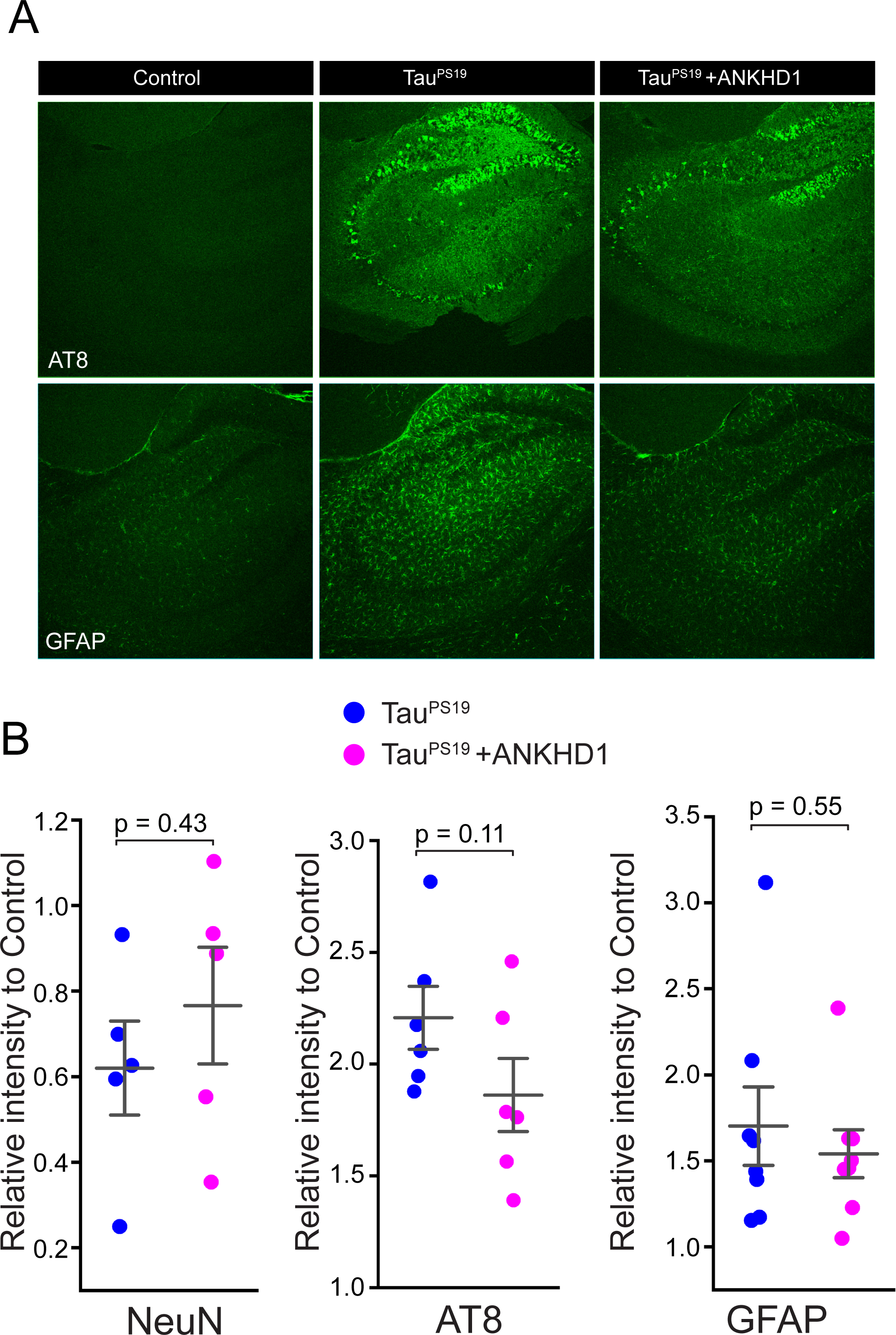
The effect of ANKHD1 co-expression on the Tau phosphorylation and gliosis in TauP301S transgenic mice at 9-month of age. (**A**) Representative confocal images of mouse brain hippocampus of Control, Tau^P301S^, and Tau^P301S^ + ANKHD1 mice. Mouse brain sections were stained with AT8 (anti-Tau-phospho-Ser202 and phosho-Thr205) and anti-GFAP antibodies, respectively, to detect hyperphosphorylated Tau and microglial cells. (**B**) NeuN, AT8, and GFAP intensities in the brain sections from male mice were quantified. Average fluorescence intensity of NeuN, AT8, or anti-GFAP was normalized to their control littermates. The data were presented as the relative intensity ratio.

### ANKHD1 partially restores cognitive functions in PS19 mice

Tau pathology in the brain causes functional abnormalities, and previous analysis of the PS19 mice by other groups showed that these mice exhibit cognitive defects at 9-month-old age when examined with novel object recognition (NOR) ^35^ and Y-Maze^34^ behavioral tests. We next tested whether the suppression of neuropathology in PS19 by ANKHD1 may correlate to improved cognitive functions. PS19 female mice showed significant drop in NOR index compared to control mice (44 vs 59). We found that the NOR index is increased from 44 in the PS19 mice to 58.5 in the PS19-ANKHD1 mice (Fig. 4). However, the reduction of NOR in PS19 male mice was not significant. Nevertheless, PS19-ANKHD1 male mice showed a NOR index indistinguishable to the control mice (Fig. 4). In our Y-Maze test, neither male nor female 9- month-old PS19 mice showed detectable deficit, precluding us to test whether the PS19- ANKHD1 mice could suppress the Tau-induced behavioral impairment in this test (Supplemental Fig. 2). These findings imply that ANKHD1’s partial restoration of cognitive function in the PS19 brain may be due to its role in reducing Tau pathology, thus highlighting its potential as a therapeutic target.

**Figure 4.**
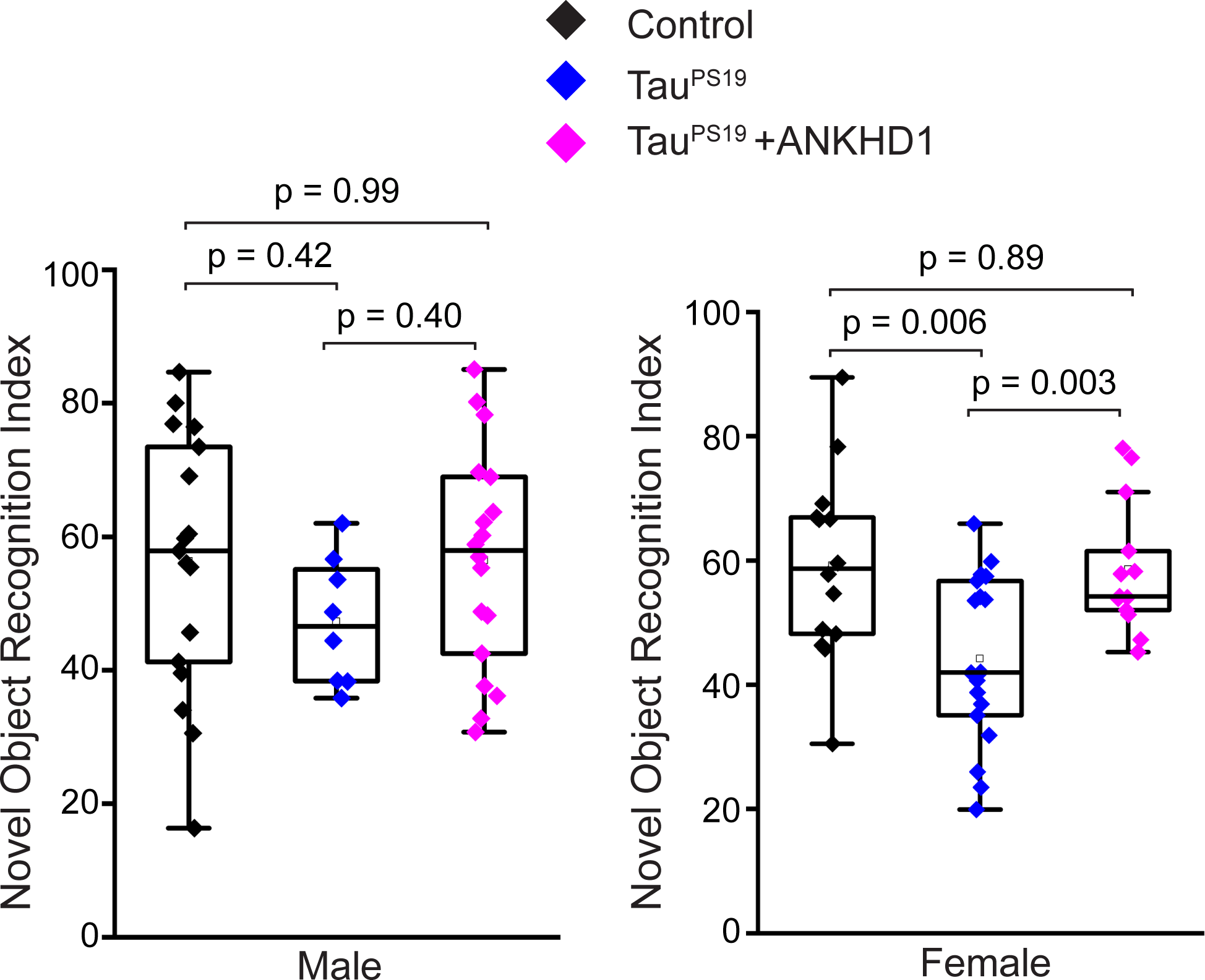
ANKHD1 co-expression rescues the defective NOR in TauP301S transgenic female mice at 9-month of age. Control, Tau^P301S^, and Tau^P301S^ + ANKHD1 mice were tested for their ability to recognize novel objects following standard protocol. Novel Object Recognition Index from the male and the female cohorts were presented.

## Discussion

We previously reported that overexpressing Mask, the fly homolog of mammalian ANKHD1 and ANKRD17, mitigates neuronal degeneration caused by human Tau and FUS in the fly eyes^18^.

Our findings reveal that Mask boosts autophagic flux, which is essential for its protective role against conditions that lead to neuronal degeneration and death. Mask and ANKHD1 exhibit strong structural conservation, suggesting they share functional similarity. Here we present evidence to support the notion that ANKHD1, like its fly counterpart, promotes autophagy in cells and ameliorates neuropathology in the mouse models for AD. ANKHD1’s role in promoting autophagy may directly counteract Tau pathology by enhancing the clearance of toxic Tau aggregates. Autophagy is crucial for degrading misfolded proteins and aggregates, including hyperphosphorylated Tau, which is central to neurotoxicity in tauopathies. By upregulating ANKHD1, we observed a decrease in p62 levels, suggesting enhanced autophagic flux, which likely facilitates the removal of hyperphosphorylated Tau. This reduction in Tau load aligns with the observed mitigation of cognitive deficits in TauPS19 mice, indicating that ANKHD1- mediated autophagy not only addresses cellular pathology but also translates into functional recovery.

We showed that increasing ANKHD1 expression in HEK293 cells boosts autophagy level, mirroring the effect of Mask upregulation in fly larval muscles. We next induced ANKHD1 expression in the brain of PS19 mouse and examined whether co-expressing ANKHD1 with human mutant Tau suppresses the AD-linked degeneration. Our results demonstrated that ANKHD1 mitigated the accumulation of the hyperphosphorylated Tau protein in hippocampus (Fig. 3, 4) as well as the related impairment of cognitive functions (Fig. 5). However, compared to the prominent protective effects of Mask in the fly eyes, ANKHD1 only induces moderate effects in mice. What accounts for the weaker protective effect of ANKHD1 against Tau-induced neurodegeneration in mice compared to its counterpart in flies? This is unlikely attributable to possible lesions on the ANKHD1 transgene elements, as all DNA sequences from the promoter to the 3’ end of the ANKHD1 are intact in the transgenic lines (supplemental Fig. 3). We speculated that one potential reason is that the neuronal population of CamKII-Cre-positive neurons does not fully align with the population expressing human mutant Tau driven by the prion promoter in the PS19 mice. Since CamKII-Cre is utilized to induce ANKHD1 expression, the ANKHD1-expressing neurons do not completely overlap with those expressing mutant Tau protein. Further studies using the same promoter to drive the ANKHD1 and human mutated Tau expression will test whether ANKHD1 acts as a potent neuronal protector like its counterpart in flies.

Furthermore, Mask and ANKHD1 share significant sequence homology, particularly in the ankyrin repeat domains and KH domain, suggesting a conserved role in promoting autophagy. Both proteins appear to enhance autophagic flux, yet the mechanisms may differ slightly due to species-specific interactions with autophagy-related pathways. In addition, Mask is known to regulate microtubule stability in *Drosophila* neurons, impacting synaptic morphology and functions, and possibly supporting axonal transport, which is crucial in mitigating Tau toxicity. Whether ANKHD1 retains this microtubule-regulating function in mammals remains to be determined, though its neuroprotective effect against Tau pathology, as seen in the present study, implies a potentially broader role.

Future studies could further elucidate the mechanistic pathways by which ANKHD1 promotes autophagy, including its specific roles in autophagosome formation and fusion with lysosomes. Additionally, investigating whether ANKHD1 interacts with other tau clearance mechanisms or contributes to microtubule stability and axonal transportation may provide insights into its broader neuroprotective effects. Therapeutically, developing ANKHD1 agonists or mimetics and exploring gene therapy applications may offer promising approaches for tauopathy treatment.

Expanding the study of ANKHD1’s effects to other tauopathies, such as frontotemporal dementia, could determine its generalizability across neurodegenerative diseases. Finally, long-term studies are essential to assess the safety and efficacy of ANKHD1 modulation, ensuring that its benefits are neuron-specific and devoid of systemic side effects.

## Materials and Methods

### Generation of Cre-inducible ANKHD1 transgenic mouse

To generate the CreSTOPGFP-ANKHD1-IRES-mCherry DNA construct, we first PCR the IRES sequence from the pCall-IRES-GFP plasmid (a gift from Dr. Hang Shi) and cloned this sequence into the Cre Stoplight 2.4 (Addgene item #37402) in between the LoxP-GFP-STOP cassette and the coding sequence for mCherry. The resulting plasmid was then used as the vector to receive the insert of the ANKHD1 coding sequence that was cut out from the Mask1 plasmid (a gift from Dr. xxxx and ref). The final DNA construct was sent to Texas A&M Institute for Genomic Medicine for perinuclear injection. C57/B6 strain was used for the injection for generating the founder lines. ANKHD1 internal sequence at the 3’ end of the coding sequence was used for PCR genotyping (Forward primer: GCACGTGGGCACCTCATATT; and Reverse primer: GGTACCTTCTGGGCATCCTT), Myogenin locus was used as the control for genotyping (Forward primer: TTACGTCCATCGTGGACAGC; and Reverse primer: TGGGCTGGGTGTTAGCCTTA). Other primers used for confirming the integration of the transgene are: TransgeneCMVFor: CCATGGTGATGCGGTTTTGGCA; TransgeneGFPRev: CCTCCATGCGGTACTTCATGGT; TransEnhancerF1: GCCAGATATACGCGTTGACATT; TrsnsEnhancerR1: GCTATCCACGCCCATTGATGTA.

### Cell culture and transfection

HEK293 cells were cultured in 1XDMEM (Gibco 11966-092) supplemented with 10% FBS, 1XNEAA, 110mg/L Sodium Pyruvate, and 100U/ml Pen-Strep antibiotics mix under optimal conditions (37°C, 5% CO2). Cells were transfected at 80% confluency with CreSTOPGFP- ANKHD1-IRES-mCherry plasmid alone or together with CMV-Cre plasmid using Lipofectamine3000 transfection reagent (Thermal Fisher Cat. No. L3000015) following protocol recommended by the manufacturer.

### Western Blot Analysis

Transfected HEK293 cells in a 12-well plate were harvested 48hrs after transfection. The cells were quickly rinsed with 1XPBS first and then incubated with SDS-Urea lysis buffer (9:2:1 of 100mM NaCl, 1mM EDTA, 100mM Tris-HCl pH8.0 Buffer : 10% SDS : 8M Urea) in the wells (400µl lysis buffer/well for a 12-well plate). The lysate was then treated with DTT (0.1M), mixed with 4XSDS loading buffer, boiled for 5 minutes and centrifuged at 15,000g for 5 minutes. The supernatant containing proteins was loaded and resolved on a 4-20% gradient gel (BioRad 17000436). The following primary antibodies were used for immunoblotting: rabbit anti-ANKHD1 (1:1,000, Biorbyte orb374007), rabbit anti-P62 (1:1,000, a gift from Dr. Sheng Zhang), mouse anti-GAPDH (1:10,000, Abcam, Ab181602). All HRP-conjugated secondary antibodies (from Jackson ImmunoResearch) were used at 1:5,000 dilution. Data was collected using a ChemiDoc MP Image System (Bio-Rad) and quantified using ImageQuant TL software (GE).

### Immunohistochemistry

Mice brains were dissected at appropriate ages, fixed and embedded as previously described^36^. All mice were deeply anesthetized with 5% isoflurane, and the whole brains were dissected, rinsed with cold 1XPBS and then fixed in 4 ml of 4%PFA in 1XPBS at 4°C O/N and then incubated in 10ml of 30% sucrose in 1XPBS at 4°C till the brains sink to the bottoms of the tube. The fixed brains were dried with Kim Wipe tissues and imbedded in OCT, freeze and stored at - 80°C till they were sectioned.

The brain blocks were moved to -20°C for one hour before sectioning. 20um sections were collected encompassing the hippocampus into each well of 12-well plates filled with 1XPBS buffer. The free-floating sections were used for immunohistochemistry. Two consecutive sections were kept in a single well and washed with 1×PBS 3 times, followed by 0.1% Triton X- 100/1×PBS (wash buffer) for 3 times, and then incubated with primary antibodies that were diluted in the blocking buffer (1XPBS containing 0.1% Triton X, 2% goat serum, and 2% BSA) at 4°C O/N. The following day, sections were rinsed in wash buffer for 3 times and then incubated with fluorophore conjugated secondary antibodies for 2 h at room temperature. The sections were placed on the slides, mounted in SlowFade Gold antifade reagent (Invitrogen S36936) and covered with coverslips for confocal microscope observation. Primary antibodies used in this study were as follows: mouse anti-phosphos-Tau Ser-202, Thr205 (1:100, AT8- biotin, Thermo Scientific MN1020B), rabbit anti-GFAP (1:1,000, Cell Signaling #3670), mouse anti-NeuN (1:500, Millipore Sigma MAB377), rabbit anti-NeuN (1:1,000, Cell Signaling #24307), and rabbit anti-Iba-1 (1:1,000, Cell Signaling #17198). For secondary antibodies, 488-, Cy3- or Cy5-conjugated donkey anti-species antibodies were used (all 1:1,000, Jackson ImmunoResearch).

### Confocal microscopy and quantification

Single-layer or z-stack confocal images were captured on a Nikon (Tokyo, Japan) C1 confocal microscope. Images shown in the same figure were acquired using the same gain from samples that had been simultaneously fixed and stained. The mean intensities of NeuN, GFAP, and phosphorylated Tau (AT8) were quantified using NIS-Elements imaging software (Nikon).

### Y-Maze

This test is based on the willingness of rodents to explore a new environment (Ann-Katrin Kraeuter et al, 2019). Normal rodents will prefer to experience a different arm of the maze than the one they visited on their previous entry. Following previously established protocols^36^, the mice were tested in a three-arm maze containing 3 equal arms of 35 cm length, 7 cm width, and 15 cm height, attached at 120 degree angles. Each arm of the maze is labeled as arm A, B, or C. In each session, the animal is placed in the center and allowed to freely explore the three arms for 8 minutes. Number of arm entries and number of alternations are scored live in order to calculate the percent alternation. An entry is considered as occurred when all four limbs are within the arm. The alternation percentage is calculated by dividing the number of alternations by number of possible triads x 100. The maze is cleaned with Chlorhexidine solution between animals to eliminate odor traces. All videos were recorded with and analyzed by Any-maze Y-Maze protocol (Any-maze).

### Novel Object Recognition

Novel Object Recognition (NOR) task was performed as described previously^36^. The method is based on the observation that rodents tend to spend more time exploring a novel object than one they’ve previously encountered. For object recognition, animals were first placed in the square chamber allowing free exploration for 10 minutes to habituate the environment. One hour later, the animals were put back in the chamber containing two identical objects, and the animal was given 5 minutes to explore. Another hour after this training, the animal was returned to the chamber, and this time, one object was replaced with a novel object that had not been previously encountered, and the animal was allowed to explore for 5 minutes. A video camera recorded movement in the chamber, and the time spent exploring each object was scored by Any-Maze NOR. The time spent exploring the novel object relative to the familiar one served as a measure of memory: the relative exploration time was recorded and expressed by a discrimination index (DI = (tnovel − tfamiliar)/(tnovel + tfamiliar) × 100%). To eliminate any olfactory cues, the chamber and objects were thoroughly cleaned after each use.

### Statistical analysis

Statistical analysis was performed, and graphs were generated using OriginPro (OriginLab, Northampton, MA). Each data set was tested for normal distribution and then compared with other samples in the group (more than two) using one-way ANOVA followed by post-hoc analysis with Tukey’s test, or with the other sample in a group of two using t-test. All Bar Overlap diagrams show mean±s.e.m with all data points indicated in each graph.

## Acknowledgements

This work is supported by a NIH/NINDS grant (NS123861) to C.W. and a LIFT grant from LSU to C.W. and X.T.

## Figure Legend for the supplemental figures

**Supplemental Figure 1.**
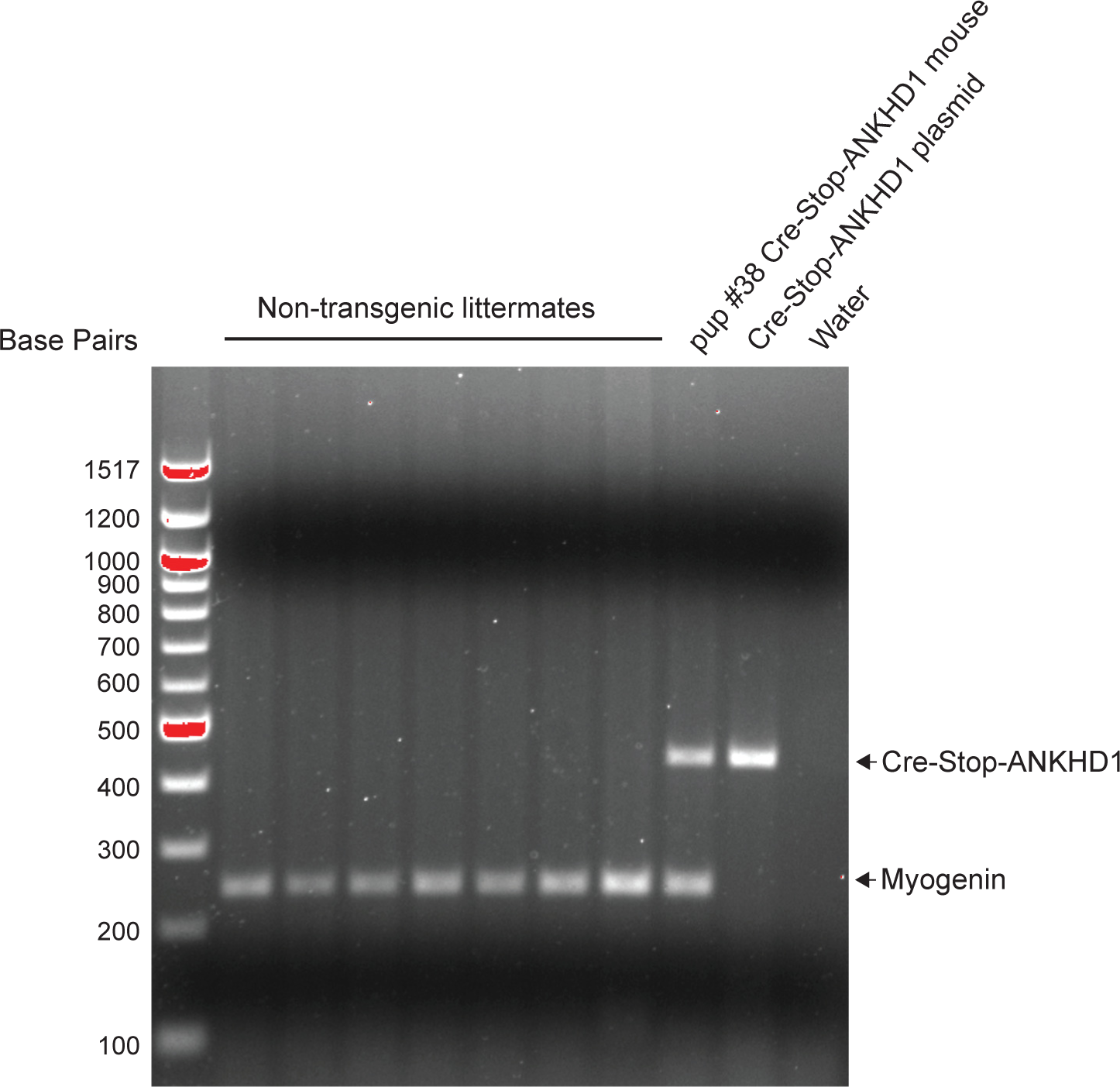
G**e**notyping **of the Cre-Stop-ANKHD1 mouse.** Genomic DNAs from the punched ear tissues of transgenic and non-transgenic pups were extracted and subjected to PCR genotyping with primer pairs recognizing *Myogenin* and the transgenic *Cre-Stop-ANKHD1.* Cre-Stop-ANKHD1 DNA plasmid was used as a positive control and water was used as a negative control.

**Supplemental Figure 2.**
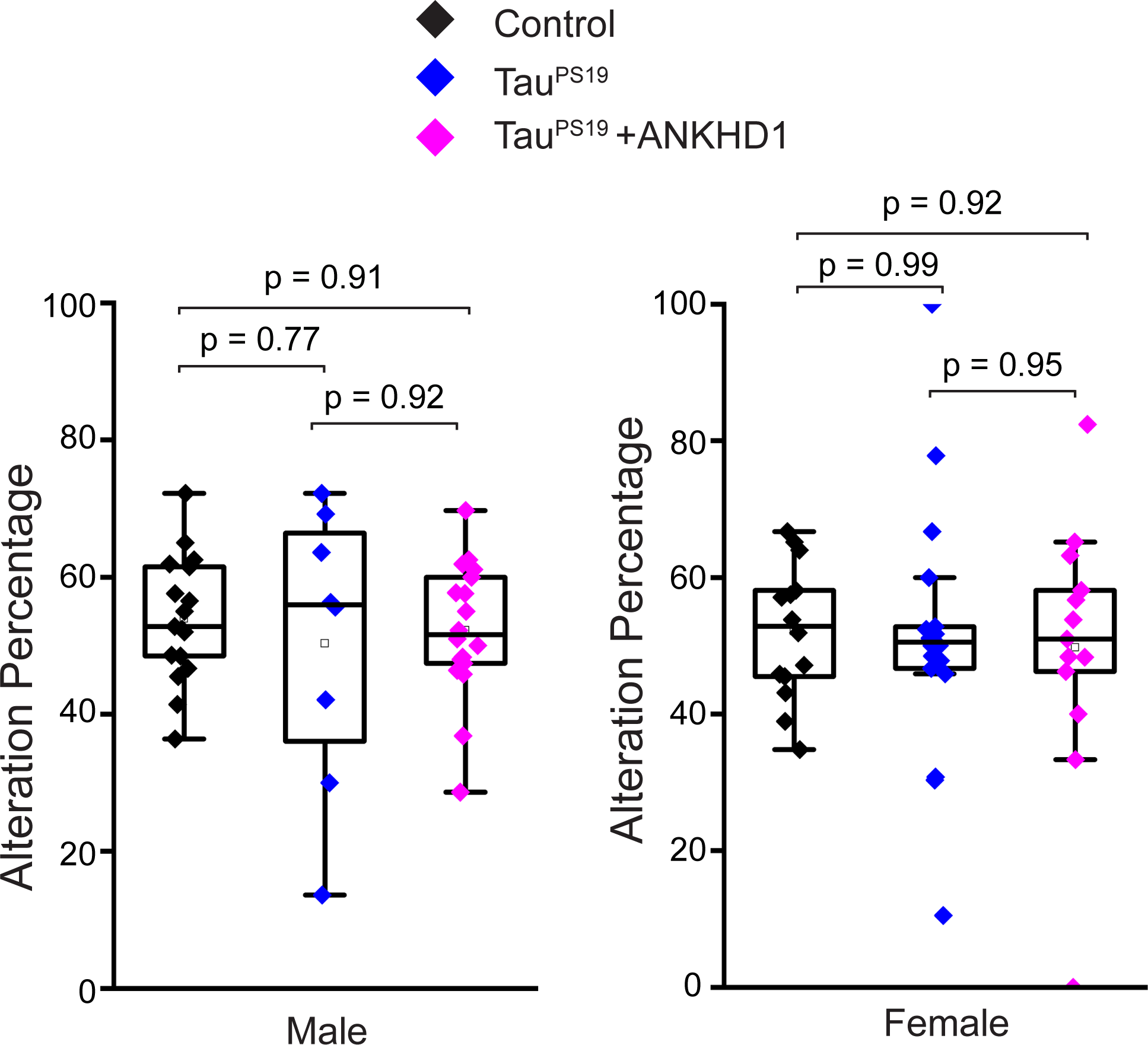
Y-Maze test of 9-month old mice. Control, Tau^P301S^, and Tau^P301S^ + ANKHD1 mice were subjected to Y-Maze test following standard protocol. Alteration Percentage from the male and the female cohorts were presented.

**Supplemental Figure 3.**
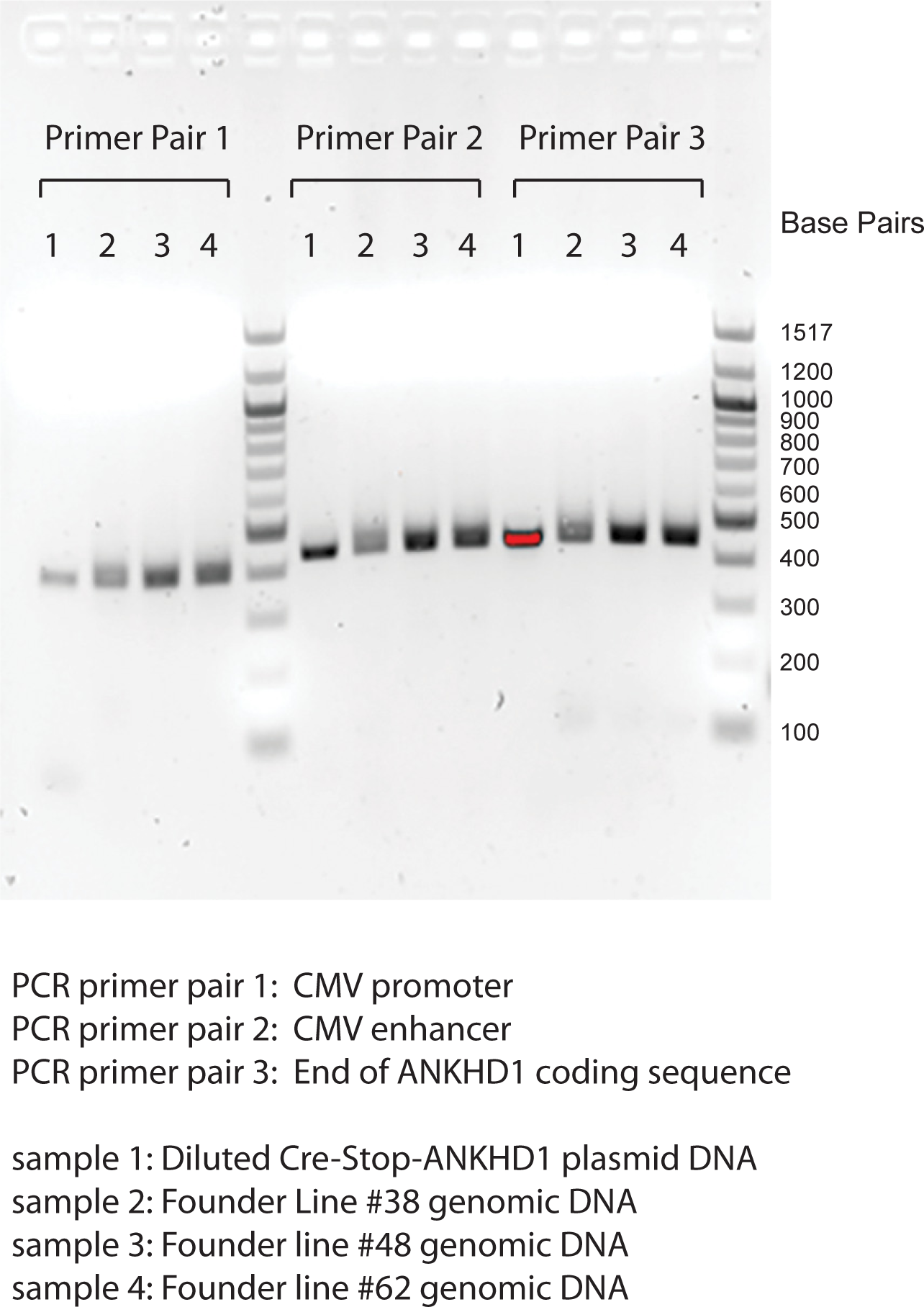
V**a**lidation **of the sequence integrity of the Cre-Stop-ANKHD1 transgene insertion.** Diluted DNAs of Cre-Stop-ANKHD1plasmid or genomic DNAs from three independent transgenic *Cre-Stop-ANKHD1*funder lines were subjected to PCR reactions with three primer pairs. The three primer pairs encompass the promoter region to the 3’ end of the ANKHD1 cDNA sequence in the *Cre-Stop-ANKHD1*transgene, as listed in the figure.

## Notes

### Competing Interest Statement

The authors have declared no competing interest.

